# Single-Cell Profiling Resolved Transcriptional Alterations and Lineage Dynamics of Subventricular Zone after Mild Traumatic Brain Injury

**DOI:** 10.1101/2021.05.31.446381

**Authors:** Xiameng Chen, Shuqiang Cao, Yinji Wang, Manrui Li, Yadong Guo, Yi Ye, Zheng Wang, Hao Dai, Wenjie Yang, Yuwen Sun, Xiaofeng Ou, Min He, Xiaoqi Xie, Weibo Liang

**Affiliations:** Department of Forensic Pathology and Forensic Clinical Medicine, West China School of Basic Medical Sciences and Forensic Medicine, Sichuan University, Chengdu, China; Department of Forensic Genetics, West China School of Basic Medical Sciences and Forensic Medicine, Sichuan University, Chengdu, China; Department of Forensic Science, School of Basic Medical Sciences, Central South University, Changsha, Hunan 410013, China; Department of Forensic Toxicological Analysis, West China School of Basic Medical Sciences & Forensic Medicine, Sichuan University, Chengdu 610041, Sichuan, China; Institute of Forensic Medicine, West China School of Basic Science and Forensic Medicine, Sichuan University, Chengdu, China; West China School of Basic Medical Sciences and Forensic Medicine, Sichuan University, Chengdu, China; Department of Critical Care Medicine, West China Hospital, Sichuan University, Chengdu, China

## Abstract

Mild traumatic brain injury (mTBI) is the most common form of brain trauma caused by physical impact. The subventricular zone (SVZ) is a neurogenetic niche that contributes to homeostasis and repair after brain injury. It is particularly challenging to fully elucidate the molecular alterations in the SVZ occurring in response to injury due to its cell diversity and the complex network. In this study, we aimed to address this issue using a novel transcriptomic technique- unbiased single-cell RNA sequencing. We resolved previous unknown cell subpopulations harbored in the niche, and uncovered cell type-specific alterations in gene expression, enriched pathways, and cell-cell crosstalk following mTBI. Notably, we also report novel lineage trajectories and molecular hallmarks that govern neurogenesis. This study dissects the delicate transcriptome changes of individual cell types as well as the reprogramming process of cells in the SVZ niche after mTBI, and our findings are expected to facilitate the development of therapeutic interventions or diagnostic tests for mTBI.

## Introduction

Mild traumatic brain injury (mTBI), “an acute brain injury resulting from mechanical injury to the head from external physical forces” (1), is the most common type of traumatic brain injury (2). A single mTBI could induce pathophysiological changes in the brain, leading to acute neuro-dysfunction and an array of subsequent post-acute or chronic sequel (3, 4), such as depression, anxiety, headache, irritability, and post-traumatic stress disorder. To date, specific and effective biomarkers of mTBI are absent, and the diagnosis or treatment for mTBI presents a challenge. Hence, understanding the pathogenesis and molecular regulation of mTBI are critical for improving therapeutic options.

The subventricular zone (SVZ) is one of the neurogenetic niches in the adult mouse brain and is responsible for maintaining tissue homeostasis is involved in recovery in response to stimulation (5). This neurogenetic niche is a complex micro-environment comprised of morphologically and molecularly distinct subtypes of neural stem cells (NSCs), giving rise to specific progenitor cell types committed to neuron or glia differentiation (6). Upon brain injury, NSCs in the SVZ reprogram to induce neurogenesis (7). However, most of the newly generated neural cells are gliocytes, and the ability of the brain to replace lost neurons is restricted (8). As such, the diversity of NSCs/progenitors and compartments that determine the neurogenesis are not totally apparent. To this end, understanding the diversity of the niche cells, as well as their dynamics upon injury, is critical for elucidating the pathogenesis of mTBI and identifying relevant diagnostic or therapeutic targets.

Previous studies have generally described the processes involved in the proliferation and differentiation of cells during neurogenesis. Upon activation, actively dividing NSCs (aNSCs) give rise to transit amplifying cells (TACs), which generate neuroblasts (NBs) that migrate and terminally differentiate into neurons (6). In spite of this, recent efforts are continuing to enrich the diversity of NSCs and progenitors harbored in the heterogeneous pool (9), whereas their lineage trajectories and the delicate intercellular communications are not well understood. Jens et al recently proposed that the striatal astrocytes were latent NSCs that triggers the genesis of neuronal lineage cells upstream of NSCs (10). However, whether niche astrocytes reprogramming was the initial stage of neurogenesis in the SVZ remains unclear. In addition, very little is known about the molecular cues that govern these recently discovered individual niche cells, especially under the conditions of injury.

In that regard, we performed high throughput single cell profiling- a powerful approach to resolve genetic information of individual cells (11, 12), to uncover and compare cellular heterogeneity and cellular dynamics of SVZ from controlled cortical impact (CCI) induced mTBI with those of controls. Specifically, we used droplet-based scRNA-seq (Drop-seq) (13) to reveal the most vulnerable genes in each cell type and their enriched pathways under mTBI, and unravel the post-mTBI alteration of NSC/progenitor-based regulatory networks, in order to provide potential targets for the diagnosis and treatment of mTBI. Additionally, we also unveiled a previously unknown reprogramming route of neurogenesis in the niche, and explained why the replacement of lost neurons is limited after mTBI via pseudotime analysis. Furthermore, we identified the molecular hallmarks which may contribute to the government of lineage priming, NSC activation/ quiescence and cell fate determination, providing potential targets for the development of regenerative medicines.

## Results

### The Entire Adult SVZ Niche at Single-Cell Profiling

To elucidate molecular features of the adult NSCs, their progeny at each differentiation stage, and surrounding niche cells in SVZ under CCI conditions, >15,000 cells were profiled from matched SVZs of 3 mice that had received CCI and 3 sham-operated controls. These individual cells were micro-dissected and suspended via an optimized protocol without cell sorting to best preserve the *in vivo* conditions. A chromium controller chip was used to load the pooled samples, barcode each cell, and prepare and sequence the mRNA library. After the elimination of doublets and red blood cells, 15,754 raw profiles were retained. Computational analysis of scRNA-seq data was performed with the Seurat package (https://satijalab.org/seurat/). PCA and UMAP plots were used to describe the cell-to-cell relationship (Fig. 1A). Subsequently, the identity of each cell cluster was annotated based on the known markers of various cell types in the SVZ.

**Fig 1.**
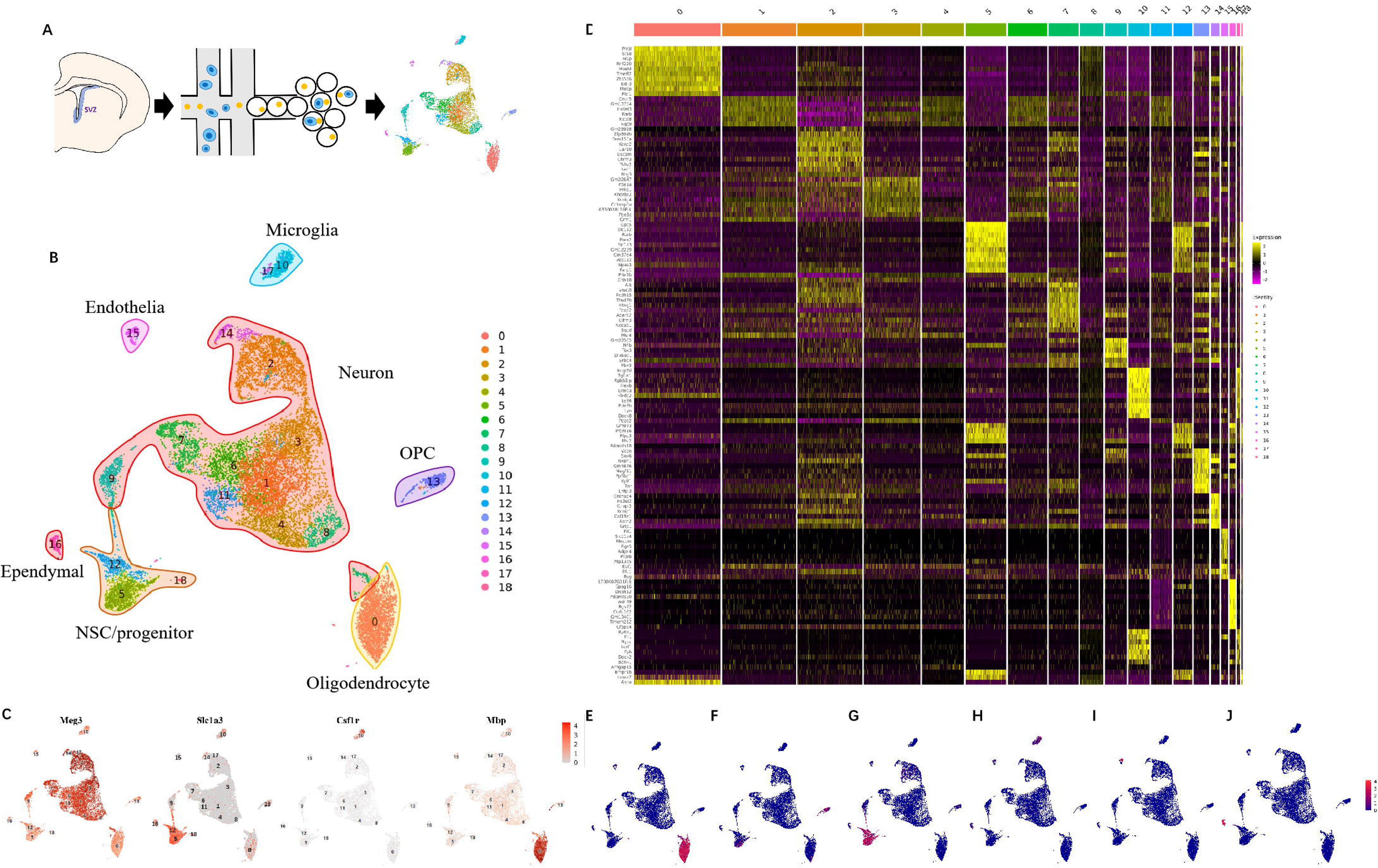
Major cell types in adult SVZ and cell type specific markers at Single-Cell Profiling. A) Workflow of sample preparation, single cell sequencing, and analysis. B) UMAP plot of single cells in the SVZ niche. C) Distribution of known markers in the UMAP: Meg3- neuron, Slc1a3- NSC/progenitor, Csf1r- Microglia, Mbp- oligodendrocyte. D) Heatmap of calculated markers in each cell cluster. E-J) Novel markers for major cell types: E- PRR5I F- KANK1, G-RORB, H- FGFBR2, I- MECOM, J- DNAH12.

In total, 18 cell clusters were detected that corresponded to NSCs/progenitors, neurons, oligodendrocyte progenitor cells (OPC) (cluster13), oligodendrocytes (cluster 0), microglia (cluster 10/17), ependymal cells (cluster 16), endothelial and mural cells (cluster 15) (Fig. 1B), based on known markers for each cell type (Fig. 1C). Since the SVZ astrocytes are usually believed to function as NSCs, and the SVZ NSCs share many similarities (including markers and molecular features) with more mature parenchymal astrocytes (7), they were preliminarily grouped together as “NSC/progenitor”. The clusters 5, 9, 12, and 18 were identified as “NSC/progenitor” according to the expression of *slc1a3* (glast), which is highly enriched in both NSCs and astrocytes (6); *atp1a2*, an astrocyte marker that is also highly expressed in NSCs (14); *egfr*, which is mostly enriched in aNSCs (7); and *aqp4*, a marker for mature astrocytes (15). Neurons were defined based on the neuronal marker *meg3* (9) and a postmitotic neuronal marker *rbfox3*. Surprisingly, ten clusters denoting various sub-types of neurons with distinct molecular features were resolved: 1, 2, 3, 4, 6, 7, 8, 9, 11, and 14 (Fig. 1D). Compared with the neuronal cell types that were previously characterized using scRNA-seq (9, 16), the current results demonstrate increased neuronal diversity in the SVZ. In addition, two distinct clusters each of microglia, ependymal cells, and oligodendrocyte lineage cells, and one cluster of endothelia and mural cells (combined as they shared the expression of some typical marker genes (16)) were detected. Furthermore, a number of novel markers for each cell type in the SVZ were identified using Drop-Seq, including *PRR5I* for oligodendrocytes, *KANK1* for OPC, *RORB* for NSC/progenitor, *FGFBR2* for microglia, *MECOM* for endothelial and mural cells, and *DNAH12* for ependymal cells (Fig. 1E-J).

### Featured transcriptome alteration in each cell type of SVZ under mTBI

Each cell type has a unique response to mTBI (Fig. 2A-I). Given that the study of cell-type specific molecular alteration could potentially reveal mTBI pathogenesis and offer therapeutic targets, the DEGs and pathways affected by mTBI were investigated based on the regional gene set defined above. *Plp1, Aldh1a1, Fry, Tac1*, and *Wdr17* exhibited the largest fold change in the NSC/progenitor (Fig. 2C); however, their effects in NSC/progenitor have never been reported. *Plp1* is an oligodendrocyte identity gene, and the large increase in *Plp1* in NSC/progenitor (Fig. 2C) indicates an oligodendrocyte-lineage differentiation in mTBI. The *Aldh1a1* gene, previously reported as a target of cancer stem cells (17), was also upregulated in NSC/progenitor post-mTBI (Fig. 2C). Its function in mTBI remains unknown, but it is worth exploring whether *Aldh1a1* could serve as a novel target for the manipulation of the NSC/progenitor niche. Studies on the *Fry* gene in the central nervous system are limited. GO annotations revealed that *Fry* could be a regulator of cell differentiation; hence, the increased *Fry* expression in NSC/progenitor (Fig. 2C) indicated a possible role for Fry in NSC/progenitor fate determination following mTBI. *Tac1* might be involved in the processes of cell death or proliferation; the significant decrease in *Tac1* (Fig. 2C) suggests that it may serve as a potential therapeutic target for mTBI. *Wdr17* has scarcely been studied, and its function remains largely unknown; thus, the greatly reduced *Wdr17* expression in NSC/progenitor (Fig. 2C) needs further investigation. Furthermore, quantitative set analysis for gene expression (QuSAGE) 2.16.1 (18) analysis revealed several DEG-enriched pathways in NSC/progenitor responding to mTBI (Fig. S1) For instance, the ferroptosis (19) pathway, a newly discovered type of programmed cell death, is specifically activated in NSC/progenitor post mTBI (Fig. S1). Ferroptosis has been reported to participate in the progression of neurodegenerative diseases and neuronal death following mTBI (20); However, very few studies have reported on SVZ NSC/progenitor-specific ferroptosis events post-mTBI. The enhanced ferroptosis in NSC/progenitor may be critical for the renewal of neural cells after mTBI and might contribute to the limited replacement of neurons after brain injury. Moreover, classical pathways related to cell proliferation or differentiation, including Ras/Rap1/PI3K-Akt signaling pathways, were found to be activated in NSC/progenitor after mTBI (Fig. 2D).

**Fig 2.**
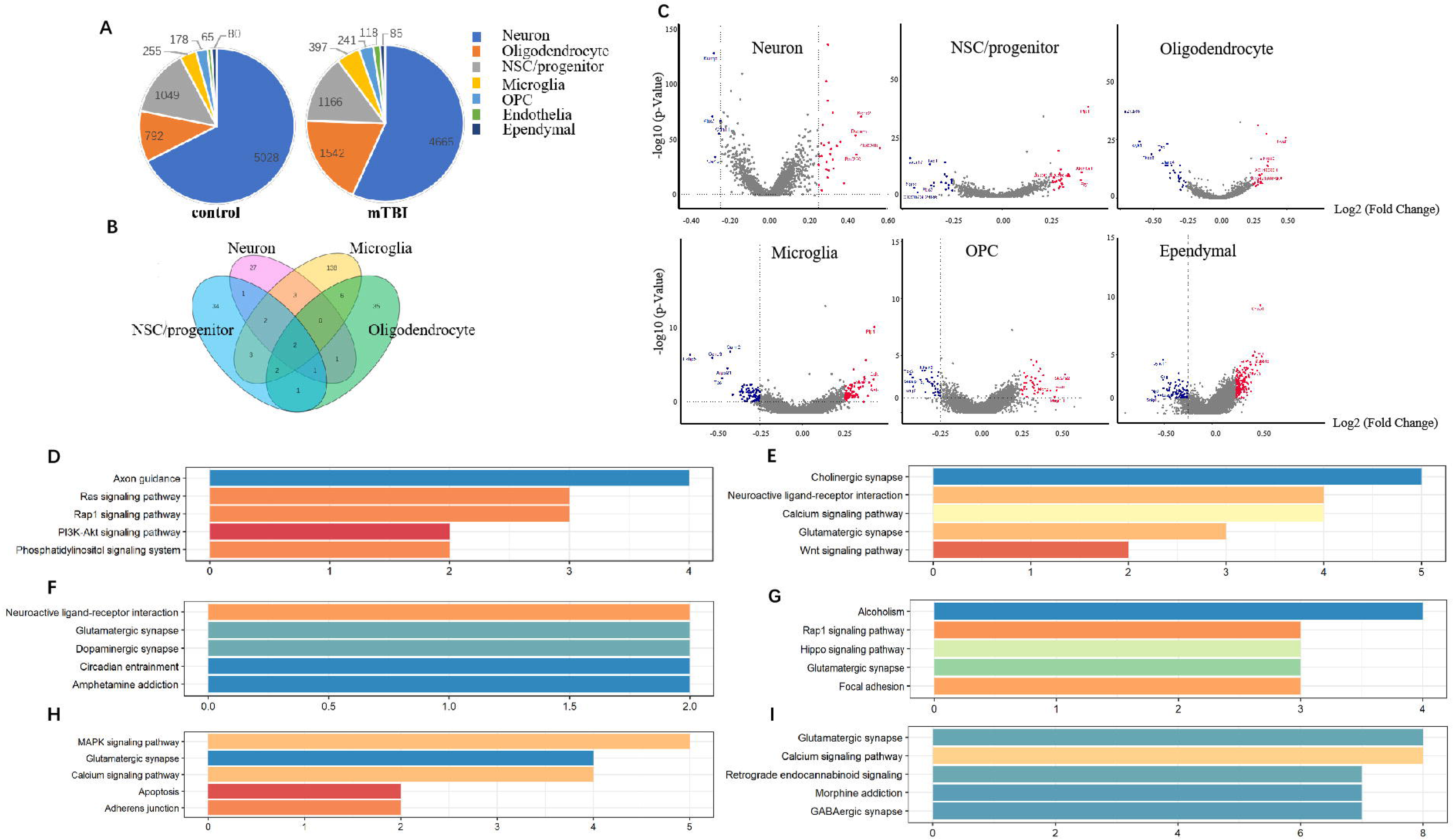
mTBI induced featured transcriptional alterations in each cell type. A) Proportions of each cell type in the mTBI and the control SVZ. B) Venn diagram showing overlap of DEGs among major cell types. C) Comparing gene expression mTBI versus control in major cell types via volcano plot. D-I) DEGs-enriched pathways in each cell type responding to mTBI. D- NSC/progenitor, E- neuron, F- oligodendrocyte, G- OPC, H-microglia, I-ependymal cell.

In neurons, the most vulnerable genes showing a response to mTBI include *Zfp804b, Kcnc2, Rnf220, Dscam*, and *Ctnna3* (Fig. 2C). Pathways enriched with these genes are related to cholinergic synapse, neuroactive ligand- receptor interaction, calcium signaling pathway, glutamatergic synapse, and Wnt signaling pathway (Fig. 2E). For other major cell types, the expression of *Neat1, Tac1, Fkbp5, Sgk1*, and *Zbtb16* in oligodendrocytes (Fig. 2C), *Slc24a2, Eml1, Magi1, Mobp*, and *Tox3* in OPCs (Fig. 2C), *Plp1, Arpp21, Tac1, Ccnd3*, and *Fkbp5* in microglia (Fig. 2C), and *Cmss1, Txnip, Sntg1, Dlg2*, and *Tubb4b* in ependymal cells (Fig. 2C) were significantly changed in the post-mTBI SVZs. Pathways enriched with these genes mainly include neuroactive ligand-receptor interaction, glutamatergic synapse, Rap1/Hippo/MAPK signaling pathways, and Calcium signaling pathways (Fig. 2F-I). In summary, the present study identified the unique DEGs and their detailed functions in each cell type of the SVZ, providing novel insights into mTBI pathogenesis and offering new therapeutic targets for mTBI that the bulk-tissue RNA-seq study may omit.

### Construction of an NSC/progenitor-based regulatory network for mTBI

In addition to the gene expression in individual cells, cellular function and behavior can also be influenced by their interaction patterns. The cell-cell crosstalk among different cell types defined above was investigated using CellphoneDB2 (Fig. 3A). It was observed that most of the gene co-expression patterns were changed post mTBI (Fig. 3A). For instance, the interactions between NSC progenitor and endothelia/mural, and ependymal cells were enhanced, whereas the crosstalk between OPCs and NSC/progenitor was subdued (Fig. 3A). Here, we focused on NSC/progenitor-based interaction networks, considering their critical role in neural cell renewal. Detailed changes in ligand-receptor association are presented in a bubble plot (Fig. 3B, C). As a receptor, NSC/progenitor interacted with a number of cell proliferation-, differentiation-, or death-related signals. For example, the NSC/progenitor *Egfr* showed an especially strong correlation with *Nrg1* from ependymal cells post mTBI (Fig. 3B). *Egfr* is known to affect cell proliferation, differentiation, migration, and survival (21, 22), which may indicate that a strategy to control NSC/progenitor behavior may include ependymal-NSC/progenitor crosstalk through the *Nrg1-Egfr* pathway; however, these results require further validation. Other targetable receptors of NSC/progenitor mainly belong to the Fgfr, Eph, Notch, and Pdgfr pathways. Under mTBI insult, the NSC/progenitor also act as source cells to perturb signaling in other cell types, most of which affect the immune system, response to stimulus, cellular component organization, and cell differentiation. Taken together, the breadth of our study enables us to disclose the elaborate alterations of molecular network after mTBI, pinpointing potential targets for its treatment or diagnosis.

**Fig 3.**
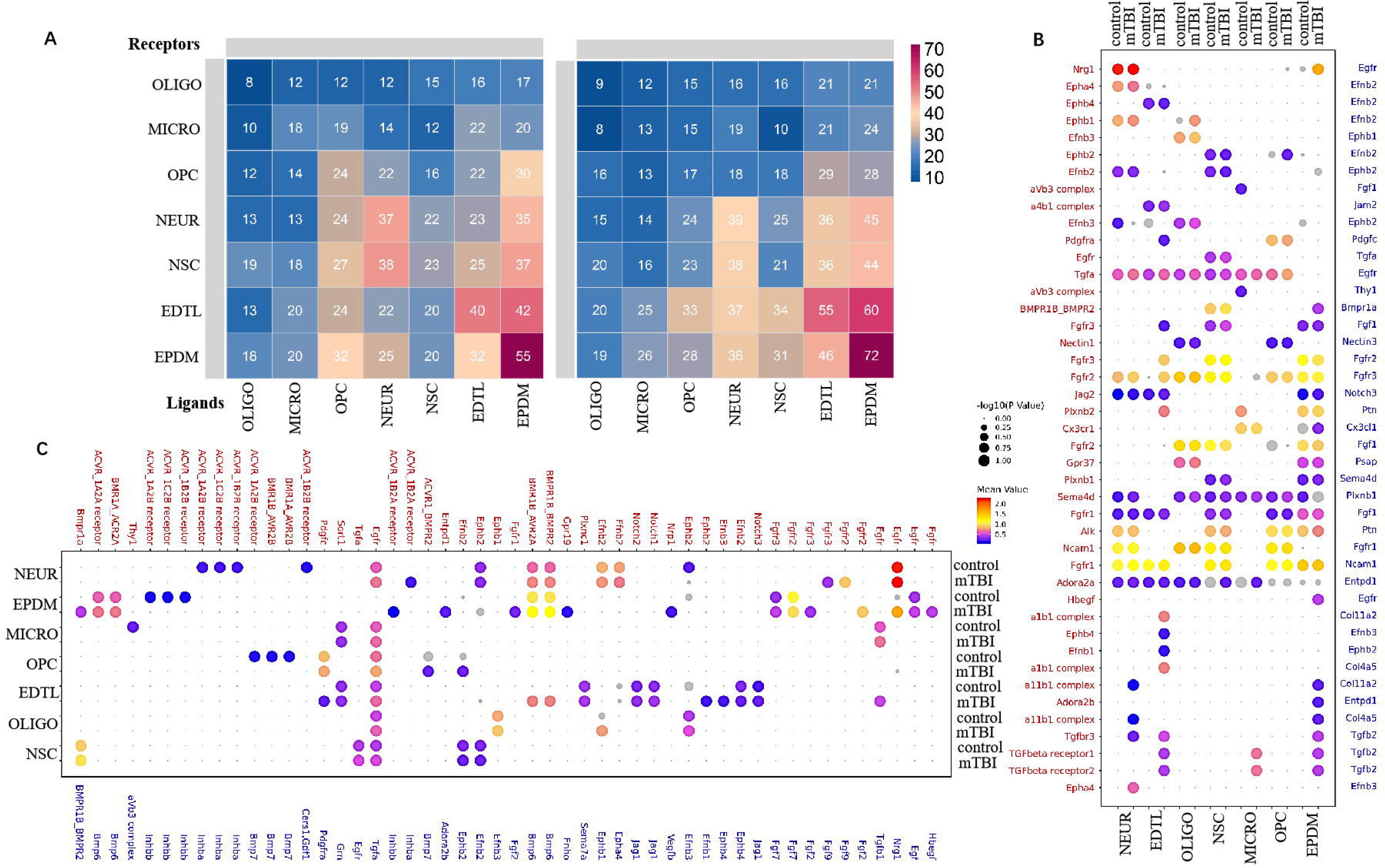
Cell to cell crosstalk was changed under mTBI. A) CellPhoneDB was applied to calculate cell-cell communication in the mTBI and control SVZ. B-C) Bubble plot showing detailed alteration of ligand-receptor interaction between cells. B: NSC/progenitor as receptor, C: NSC/progenitor as ligands. Blue font- ligands, Red font- receptors. OLIGO: oligodendrocyte, MICRO: microglia, OPC: oligodendrocyte precursor cell, NEUR: neuron, NSC: NSC, progenitor, and astrocyte, EDTL: endothelia cell, EPDM: ependymal cell.

### Approaching NSC/progenitor diversities in the mTBI and the control SVZ

For a more detailed characterization of NSC heterogeneity within the SVZ, we further resolved the original “NSC/progenitor” cell type into 9 sub-clusters (Fig. 4A) based on distinct transcriptome parameters. The following markers from previous studies were weighed to annotate NSC subtypes. 1. *Dcx* and *meg3*, associated with neuronal commitment (7), are more enriched in cluster 3 and cluster 6 (Fig. 4B); thus, they are regarded as neuroblasts. 2. *Egfr* is expressed in aNSC, but not in quiescent NSC (qNSC)(7); therefore, cluster 4 and cluster 5 were defined as aNSCs (Fig. 4C). 3. *Cdk1*, detected in cluster 5 (Fig. 4D), is a late aNSC marker (23). In addition, cluster 5 also express a certain amount of *Dcx* (Fig. 4B), indicating a less primitive state, and was therefore identified as late aNSC. 4. Cluster 4 was determined to comprise of early aNSCs as its *Egfr* (aNSC marker) expression was at a relatively low level (Fig. 4C), and it also shared some qNSC markers, suggesting an early aNSC state. 5. The SVZ astrocytes and qNSCs shared multiple markers, which were detected in clusters 0, 1, 2, and 7. The likelihood-ratio of *Thbs4* and CD9 was used (7) to distinguish qNSCs (cluster 2, 7) from astrocytes (cluster 0, 1) (Fig. 4E). Notably, although cluster 8 expresses general NSC/progenitor markers, its molecular features do not match any currently known subtype of NSC/progenitor in the SVZ. Therefore, cluster 8 was labeled as “undefined”. On the dimension reduced image, cluster 8 was located in between astrocytes and qNSCs, indicating that it may correspond with their transition state.

**Fig 4.**
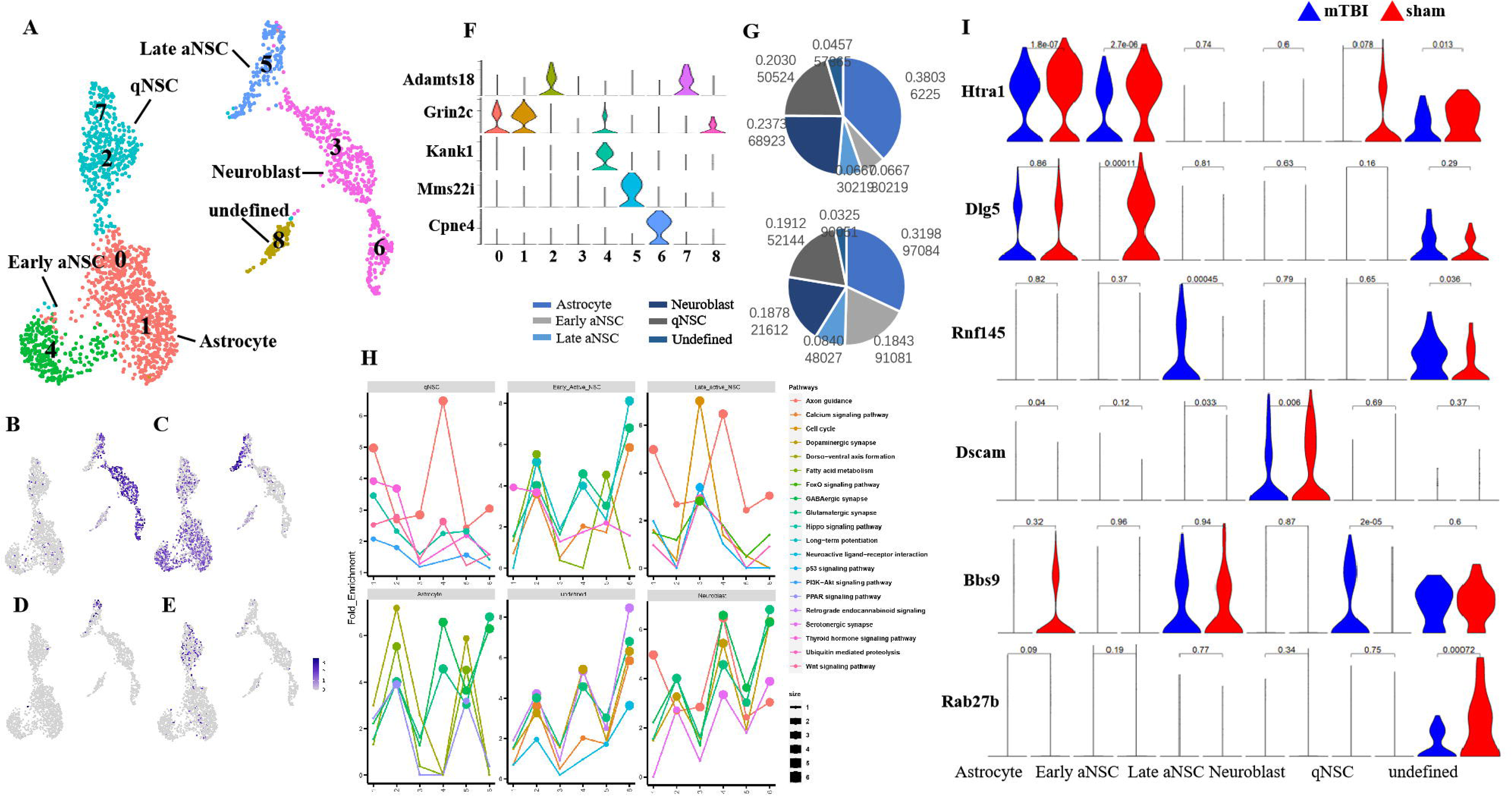
NSC/progenitor diversities in the niche and their alteration post-mTBI. A) UMAP plot showing NSC/progenitor sub-types in the SVZ niche. 9 numbers indicated different subclusters. B-F) Distribution of known markers in major sub-types: B- Dcx, C- Egfr, D- Cdk1, E- Thbs4. F) Novel markers for each sub-type. X-axis: cluster number, Y-axis: expression. G) Comparing the proportions of each sub-type in the mTBI versus the control SVZ. H) Time course of top-DEGs-enriched pathways in each sub-type of NSC/progenitor. Significance (P value) was reflected by point size. I) Violin plot showing top DEGs in each sub-type. We used Likelihood-ratio test to determine statistical significance between mTBI and control groups.

Additionally, a number of novel markers specific to NSC/progenitor subtypes in the SVZ were identified (Fig. 4F). For instance, *Adamts18* was found to be especially enriched in qNSC, but barely expressed in astrocytes (Fig. 4F); it can thus be potentially applied to distinguish these two cell types. Others like *Grin2c* in astrocytes (Fig. 4F), *Kank1* in early aNSCs (Fig. 4F), *Mms22i* in late aNSCs (Fig. 4F), and *Cpne4* in neuroblasts (Fig. 4F), also exhibited exclusive enrichment in a single subtype.

Based on the above analysis, it was observed that the proportions of qNSC and early aNSC were greatly increased within 72 h of mTBI induction (Fig. 4G), implying that the most vulnerable subtypes of NSC/progenitor respond to mTBI as well as the initial event of neurogenesis upon injury. Additionally, the comparison of the cell subtype-specific transcriptomes of the mTBI and the control SVZ uncovered a large number of DEGs and their potential functions (Fig. 4H, I), which may help normalize mTBI-induced disorders.

### Illuminating NSC/progenitor Reprogramming Trajectory in Pseudotime

To visualize the reprogramming route of NSC in SVZ under mTBI and uninjured conditions, Monocle2 was used to uncover pseudotime ordering of the NSC/progenitor cells catalogued above. Three states were detected along the route, progressing from state 1 to state 2 and state 3 (Fig. 5A). Surprisingly, we detected that the niche astrocyte (cluster 0, 1) was the major cell type that took up the tip of pseudo-time tree (Fig. 5A), indicating astrocyte reprograming as the onset of the neurogenesis process. Although there have been studies demonstrating that the niche astrocytes share many hallmarks with NSCs (7), the present study is the first to propose that the niche astrocytes triggered the neurogenic program prior to the qNSC. Subsequently, the number of niche astrocytes gradually decreased along the reprogramming route, followed by an expansion of early aNSCs and q1NSCs (cluster 2) (Fig. 5B). Upon branching, a subset of q1NSCs formed q2NSCs (cluster 7) and the undefined cell type (cluster 8), presenting a dormant state. Another group of qNSCs gradually developed into late aNSCs and then, the neuroblasts (Fig. 5A). During this period, the mTBI SVZ exhibited an expanded group of aNSCs than at the original stage, and developed an increased number of late aNSCs and less dormant qNSCs (Fig. 5B). However, the number of neuroblasts produced in mTBI SVZ was comparable to the control (Fig. 5B), suggesting that the differentiation from late aNSCs to neuroblasts is limited. This also partly explains why the adult brain has a limited ability to replace lost neurons post injury.

**Fig 5.**
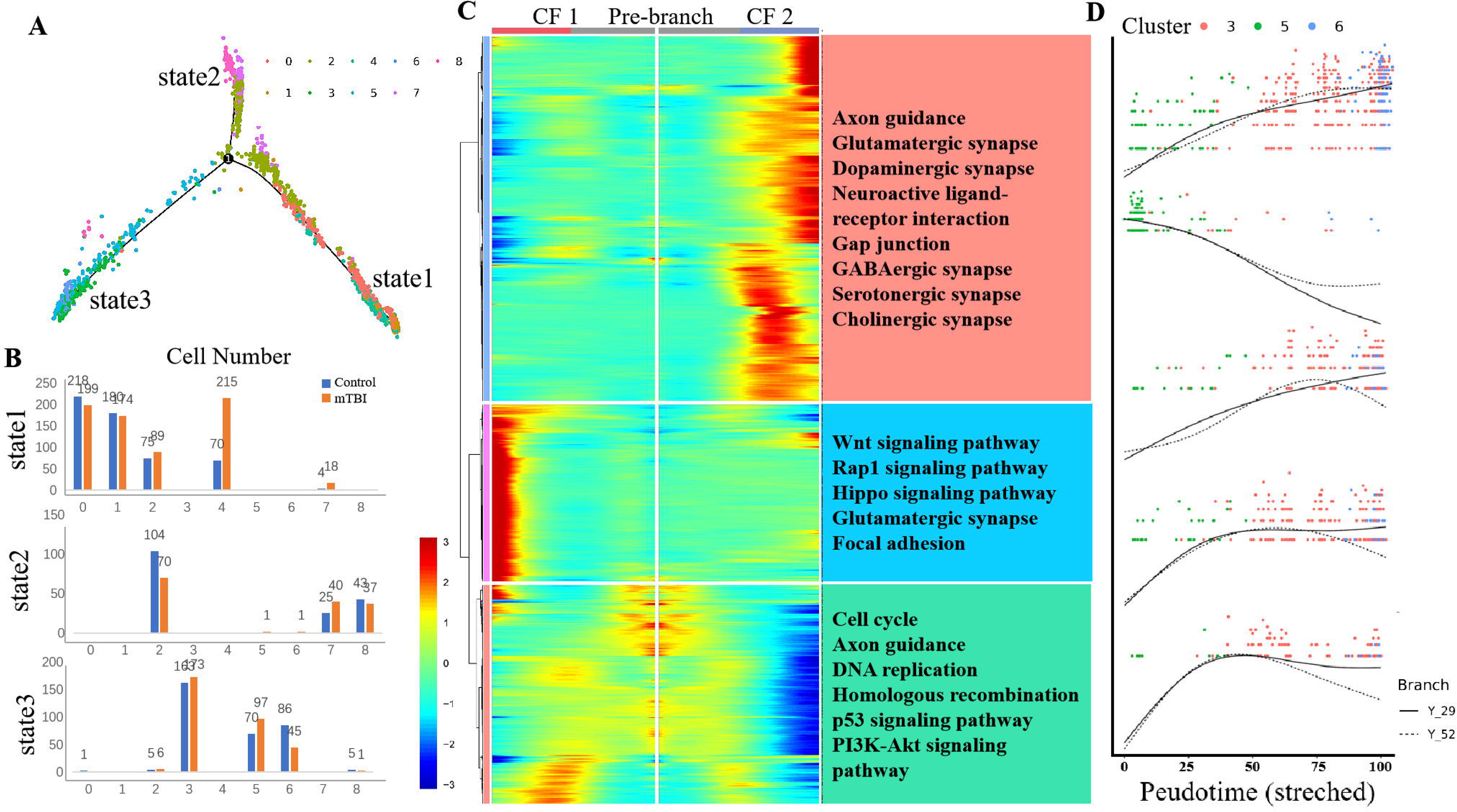
Pseudotime analysis revealed reprogramming trajectory of NSC/progenitors after injury. A) Differentiation trajectories predicted by Monocle 2, 8 colors indicated different clusters. B) Cell number comparison of each cluster at different states. C) Left: Heatmap showing gene expression patterns along the pseudotime (1000 genes). Right: 3 clusters, based on the expression dynamics, were identified. GO term for each cluster was displayed. D) Expression pattern of Cacna1c, Cdk6, Grik3, Stmn2 and Trpc4 during differentiation from aNSC to neuroblasts. Branch Y_29: cluster 3, 5; Branch Y_52: cluster 3, 5, 6.

Gene expression patterns along the reprogramming process were revealed and clustered into three clusters according to their expression dynamics (Fig. 5C). Genes responsible for NSC proliferation or differentiation as well as neuronal function altered their expression upon switching cell identity from qNSC to aNSC and neuroblasts (Fig. 5C). These included the genes involved in “Axon guidance”, “GABAergic synapse”, “Base excision repair”, “Cell cycle”, “DNA replication”, and “p53 signaling pathway” (Fig. 5C). While NSC maintenance genes, such as genes involved in “Hippo signaling pathways”, “Rap1 signaling pathways”, and “Wnt signaling pathways”, were activated in certain remaining qNSCs, the cell fate was unchanged (Fig. 5C). We further examined gene dynamics during the differentiation from aNSC to neuroblast, as neuronal supply is critical in the recovery after brain injury. The genes *Cacna1c, Cdk6, Grik3, Stmn2*, and *Trpc4* either showed a dramatic increase or a decline when aNSCs differentiated to neuroblasts (Fig. 5D). Most of these genes have not been reported to play a role in the neuron fate commitment, and deserve detailed investigation for their therapeutic potentials in post-TBI regeneration.

## Discussion

Through high-throughput single-cell profiling, we conducted an initial comparative study of the molecular biological features of each cell type/sub-type in the mTBI and control SVZ. Acknowledging the unique ability of scRNA-seq, we effectively revealed a significant amount of information about the cells, genes, and pathways vulnerable to mTBI, which may potentially provide the cellular and molecular basis for the pathogenesis of mTBI, and offer pinpointed targets for its diagnosis and treatment. While a previous study investigated the mTBI-induced transcriptional variations in the hippocampus via scRNA-seq (24), the molecular events occurring in the other neurogenetic niche, the SVZ, have not been elucidated until now. The top DEGs we detected in each cell type and the enriched pathways are distinct from those in the hippocampus (24), indicating different responses of these two neurogenic niches in mTBI. For instance, our study revealed that the top DEG-enriched pathways in SVZ neurons after mTBI include metabolic process, nutrient absorption, and cell adhesion, while Douglas et al. reported that the neurogenesis- and energy-related synaptic signaling pathways were mainly altered in the hippocampal neurons (24). Additionally, compared with those of hippocampal neurons, the diversity and function of SVZ neurons are far less clear. Our findings on the neuron diversity in the SVZ, as well as their markers and biological features, warrant future in-depth studies. In addition, we clearly show that the NSC/progenitor in neurogenic niches are critical for neurogenesis (25), which was not resolved in the previous mTBI study (24). This is the first study to report the molecular alterations in NSC/progenitors and provide an NSC/progenitor-based regulatory network for mTBI. Our study also reveals potential regulatory factors that govern the process of neurogenesis; however, further experimental validation of the data is needed.

Although intensive investigations have been performed on NSC/progenitor in neurogenesis niches (26), their identities, lineage trajectories, and molecular regulation remain debatable. By applying single-cell transcriptomics, this study resolved two subtypes of qNSCs (q1NSC and q2NSC), early aNSCs, late aNSCs, two sub-sets of niche astrocytes, two clusters of neuroblasts, and a previously unknown cell type. Notably, we uncovered a number of novel genes that may potentially mark each sub-type of NSC/progenitor, filling a gap in current knowledge. For instance, currently no specific marker effectively distinguishes qNSCs from the niche astrocytes; hence, the likelihood ratio has been applied for their identification (7). Our data detected several genes exclusively enriched in qNSCs and not in the niche astrocytes, providing potential specific markers for identification of these cell types. Another previous notion challenged by our data is that the lineage priming of neurogenesis has been determined to be initiated from the activation of qNSCs. Using pseudotime analysis, a powerful approach for revealing lineage trajectories (27, 28), we found that the niche astrocytes triggered the neurogenetic process rather than the qNSCs. Our results are consistent with a recent study suggesting that the parenchymal astrocytes initiate neurogenesis before NSC activation (10). In addition, we detected an early aNSC differentiation stage prior to the activation of qNSCs. Upon qNSC reprogramming initiates, a branch of qNSC became dormant, which contains the previously unknown sub-type, and pathway analysis also suggested the undefined cluster to be in a less active state. It will be interesting to conduct in-depth studies and further characterize these currently unknown sub-sets of NSC/progenitors. Moreover, our data provide clues for understanding the mechanisms underlying the limited replacement of lost neurons after brain injury, pinpointed the targets for solving this problem. Adding to this picture, we also clarified the molecular events orchestrating each neurogenic stage at single-cell resolution and identified potential genes that may govern the fate of each NSC/progenitor sub-type; these findings are expected to facilitate the development of regenerative medicines.

## Methods

### Experimental animals and ethical clearances

Male C57BL/6J mice, aged 8-10 weeks, were housed in cages with ad libitum access to food and water and acclimated to standard conditions (12 h light/dark cycle, temperature: 22–25 °C, relative humidity: 40-60%) for at least one week before the experiment. All experiments were approved by and conducted as per the guidelines of the Animal Care and Experimental Committee of Sichuan University, China.

### mTBI model

Each mouse received a single mild controlled cortical impact (CCI). Briefly, C57BL/6 mice were anesthetized with 4% isoflurane, with 1-2% isoflurane used subsequently for maintenance. After mounting the animal on a stereotaxic frame, a midline incision was made on the scalp of each mouse under sterile conditions. Soft tissue was carefully removed to expose the skull, and the mice were then transferred to a customed foam pad for impact. A flat tip (3.0 mm diameter) with round edge was placed 2.0 mm caudal to bregma and 3.0 mm lateral to the midline. The electromagnetic CCI device was set to deliver an impact with 4.0 m/s velocity, 1.0 mm depth, and 100 ms dwelling. Subsequently, incisions were closed with intermittent sutures followed by sterilization. After the procedure, mice were removed from the stereotaxic frame and returned to cages for recovery. Sham-operated control mice underwent the same procedures except for the impact. Animals with skull fractures, cerebral hemorrhages, or contusions were excluded from the following experiment. Mice were sacrificed 48 hours post injury, and the SVZ was micro-dissected (n=3 per group) and washed in precooled PBS to remove residual blood. Then, the tissue was flash-frozen in liquid nitrogen and transferred to −80 □ L until further use.

### scRNA-Seq

NLB buffer (X-100 (Sigma-Aldrich, St Louis, MO, USA), 0.2 U/μL RNase Inhibitor (Takara, Kyoto, Japan), 250 mM Sucrose, 10 mM Tris-HCl, 3 mM MgAc_2_, 0.1% Triton 0.1 mM EDTA) was used to homogenize the frozen tissue. Different concentrations of sucrose were applied to purify the nuclei using sucrose density gradient centrifugation. A final concentration of around 1000 nuclei/μL was used for scRNA-Seq. The scRNA-Seq libraries were built using the 10X Genomics Chromium Controller Instrument and Chromium Single Cell 3’ V3.1 Reagent Kits (10X Genomics, Pleasanton, CA, USA). Briefly, the acquired cell nuclei were loaded into several channels to obtain single-cell Gel Bead-In-Emulsions (GEMs). After reverse transcription, barcoded cDNA was released from the GEM and then purified. After amplification, the barcoded cDNA went through fragmentation, ligation with adaptors, and amplified with index PCR. The Qubit High Sensitivity DNA assay (Thermo Fisher Scientific, Waltham, MA, USA) was applied to quantify the final libraries, and the size distribution was determined by the High Sensitivity DNA chip using Bioanalyzer 2200 (Agilent Technologies, Santa Clara, CA, USA). Novaseq6000 (Illumina, San Diego, CA, USA) was used for sequencing on a 150 bp paired-end run.

### Analysis of single nucleus RNA-sequencing data

Reads from each single nucleus were aligned to the mm10 genome with Cell Ranger v3.1.0. To minimize the sample batch effect, down sample analysis was applied among samples sequenced according to the mapped barcoded reads per nucleus to finally achieve the aggregated matrix. Fastp with default parameters was used for quality control of raw reads (29). Nuclei with high levels of mitochondrially expressed genes (>20%) or containing less than 200 expressed genes were excluded from the expression table. Seurat v3.0 and various R packages were used for further analysis. Principal component analysis (PCA) was conducted using the top 2000 highly variable genes, and the uniform manifold approximation and projection (UMAP) plot was constructed based on the top 10 principals. Subsequently, the unsupervised cell clusters were defined, and marker genes for each cluster were calculated using “FindAllMarkers” with Wilcox rank sum test (logFC > 0.25; p < 0.05; min.pct>0.1). For NSC/progenitors, a semi-supervised sub-clustering was performed.

### Functional annotation and pathway Analysis

Gene ontology (GO) annotations (30) were downloaded from NCBI (http://www.ncbi.nlm.nih.gov/), Gene Ontology knowledgebase (http://www.geneontology.org/), and UniProt (http://www.uniprot.org/). Pathway analysis was performed using the KEGG database. Fisher’s exact test was applied to identify significant GO categories and pathways. FDR was used to correct the p-values, and p-values□<□0.05 were considered significant (31).

### Cell–cell gene co-expression analysis

Communications between cells were analyzed using the Python based CellPhoneDB (32) (Version 2.0), a public repository of ligands, receptors and their interactions. Interactions with p-values < 0.05 were selected for revealing relationships between cell types.

### Pseudo-time analysis

The reprogramming trajectory analysis was performed using Monocle2 (http://cole-trapnell-lab.github.io/monocle-release), with DDR-Tree and default parameters. A heatmap was produced to display the series of genes with certain expression patterns along the pseudotime. Using the differential gene test function of Monocle2, a q value <0.01 was set to identify significantly changed genes.

### Statistical analysis

All analyses were performed with GraphPad Prism (version 7.0) and R (version 3.6.0). Data were presented as means ± SD. A p-value□<□0.05 was considered statistically significant.

## Acknowledgements

N/A

---

Fig S1. Quantitative set analysis for gene expression (QuSAGE) analysis revealed DEG-enriched pathways in each cell type responding to mTBI.

