## Supplementary figures and images for "Single-Cell Profiling Resolved Transcriptional Alterations and Lineage Dynamics of Subventricular Zone after Mild Traumatic Brain Injury"

### Supplemental Figure 1

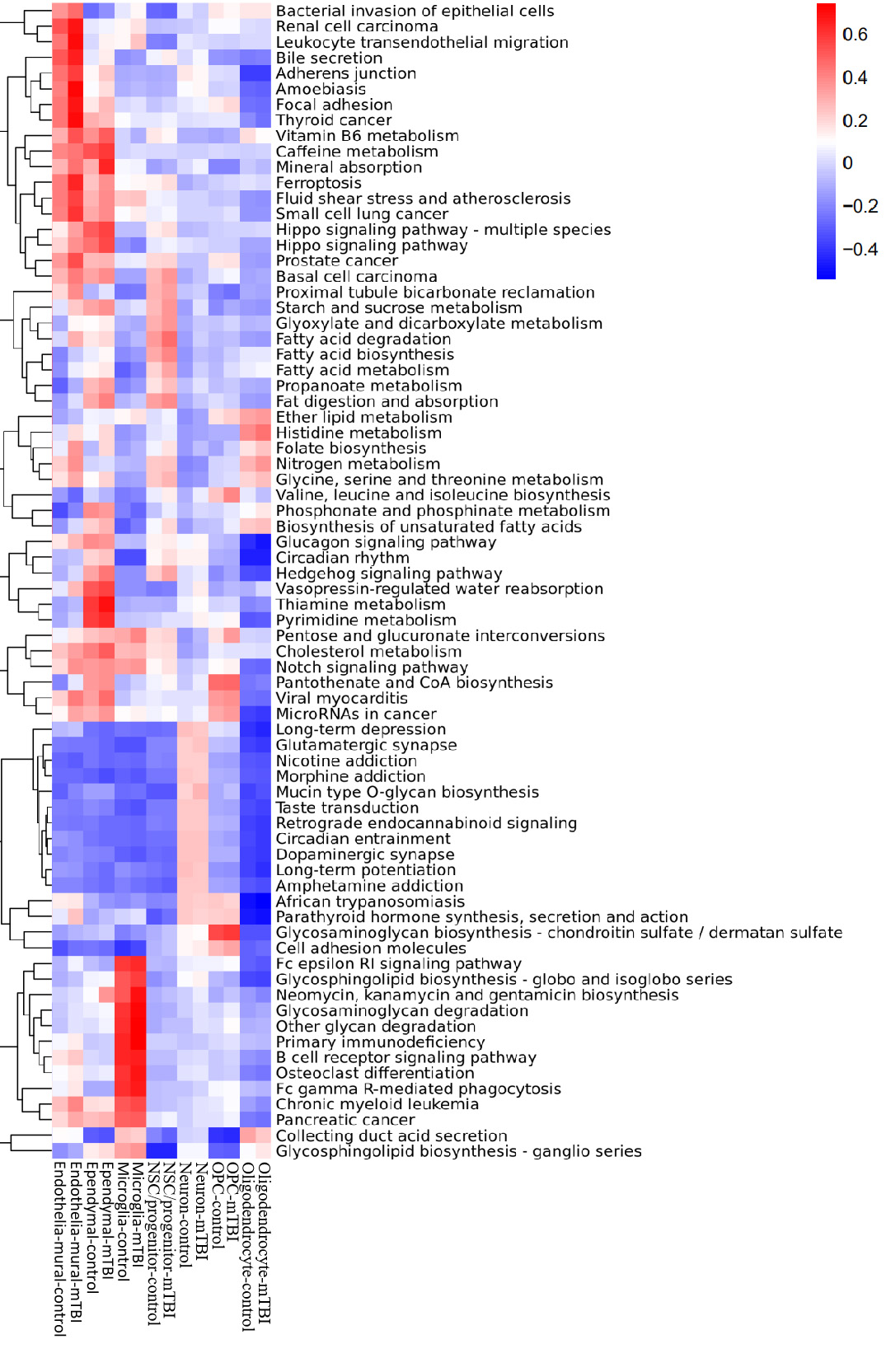
